# Automated inference of respiratory and syringeal biomechanical trajectories from birdsong acoustics

**DOI:** 10.64898/2026.08.03.742634

**Authors:** Lauren M. Ostrowski, Jorge M. Méndez, Pablo Tostado-Marcos, Brenton G. Cooper, Timothy Q. Gentner

**Affiliations:** Neurosciences Graduate Program, University of California San Diego, La Jolla, CA, 92093; Medical Scientist Training Program, University of California San Diego, La Jolla, CA, 92093; Department of Physics and Astronomy, Minnesota State University-Mankato, Mankato, MN, 56001; Department of Bioengineering, University of California San Diego, La Jolla, CA, 92093; Department of Psychology, Texas Christian University, Fort Worth, TX, 76109; Department of Psychology, University of California San Diego, La Jolla, CA, 92093; Neurobiology Section, Division of Biological Sciences, University of California San Diego, La Jolla, CA, 92093; Kavli Institute for Brain and Mind, University of California San Diego, La Jolla, CA, 92093

**Keywords:** birdsong, vocal production, biomechanical model, motor control, neural decoding

## Abstract

Songbirds, in particular zebra finches (*Taeniopygia guttata*), provide a powerful model for investigating the neural mechanisms of learned vocal behavior. Researchers typically rely on the acoustic structure of birdsong to quantify vocal behavior. As a more direct measure of motor control, we present VIBE: Vocal acoustic Inversion to Biomechanical Estimates, an open-source pipeline that recovers the biomechanical control parameters of song production directly from the acoustic waveform. Biomechanical models of the songbird syrinx describe vocal production with two continuously varying parameters: *α* and *β*, representing subsyringeal air sac pressure and syringeal muscle tension, respectively. Recovering these parameters from song acoustics provides a motor-based coordinate system against which neural activity or other dependent variables can be directly compared. Because *α* and *β* are the coupled control parameters of a nonlinear oscillator, their joint recovery is non-trivial. VIBE addresses this through iterative optimization of the governing normal-form equations. We validate VIBE against recorded air sac pressure across 44 songs from twelve birds, showing that the recovered *α* corresponds to empirically measured air sac pressure. Pairing VIBE with Neuropixels recordings from RA in five birds, we find that RA activity is well predicted by the recovered parameters, and that *α* and *β* add predictive power beyond the acoustic features of song. By recovering biomechanical control parameters from the acoustic signal, VIBE makes the biomechanical coordinate system of song production accessible to the broader songbird research community.

**New & Noteworthy:** VIBE provides a novel, fully automated pipeline to recover the biomechanical control parameters of the avian vocal organ, *α* and *β*, as continuously varying quantities from the raw acoustic waveform, making the full biomechanical model of song production accessible at the scale of modern datasets.

## Introduction

The zebra finch (*Taeniopygia guttata*) is a popular model for studying learned vocal behavior and its underlying neural activity [1, 2]. While song is measured acoustically, it is generated by motor commands issued from premotor region HVC, through the robust nucleus of the arcopallium (RA) motor region, to respiratory and syringeal vocal organ effectors [3, 4]. Thus, the acoustic features of song are an indirect readout of the motor variables under neural control. In theory, a mechanistic model of sound production can enable recovery of the biomechanical coordinates producing the song, providing a motor-relevant space against which neural activity can be directly compared [5, 6].

An established dynamical systems model of the songbird syrinx [7, 8] provides a biomechanical coordinate system of the avian vocal organ (**Fig. 1**). Each labium is modeled as a spring-mass oscillator driven by time-varying input pressure *P* (*t*); source sound passes through the trachea, treated as a tube with end reflection, then through the oropharyngeal-esophageal cavity (OEC), modeled as a Helmholtz resonator, before radiating through the beak opening [9, 10]. This model reduces to two dimensionless control parameters, *α* encoding air sac pressure and *β* encoding syringeal muscle tension, such that song becomes a smooth trajectory through (*α, β*) space [11]. Phonation onset and offset occur via a Hopf bifurcation, producing oscillations that emerge and decay smoothly. Within the phonating regime, proximity to the saddle node in a limit cycle (SNILC) bifurcation governs spectral content, with nearby trajectories yielding harmonically rich sounds and distant ones yielding tonal sounds [12, 13] (**Fig. 1**).

**Figure 1.**
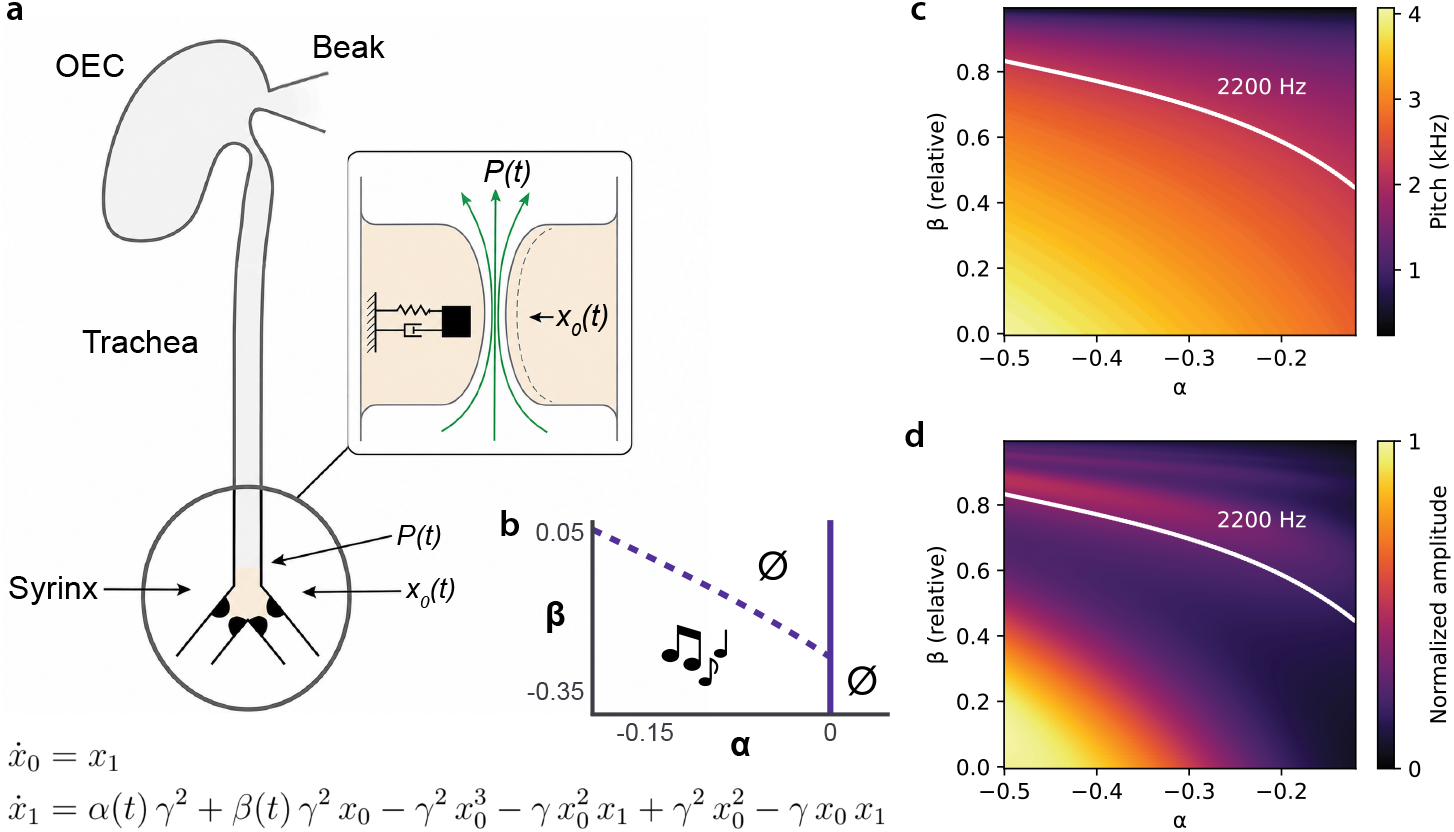
A biomechanical model of song production maps two control parameters to acoustic output. (a) Schematic of the songbird vocal apparatus, adapted from [5]. Each syringeal labium (inset) is modeled as a spring-mass oscillator with lateral displacement *x*_0_(*t*), driven by time-varying subsyringeal pressure *P* (*t*). Source sound is filtered sequentially by the trachea, modeled as a tube with end reflection, and the oropharyngeal-esophageal cavity (OEC), modeled as a Helmholtz resonator, before radiating from the beak. The governing normal-form equations are shown; *α*(t) and *β*(t) are time-varying control parameters encoding air sac pressure and syringeal muscle tension, respectively, and *γ* is a fixed timescale constant. (b) Bifurcation structure in (*α, β*) parameter space. The vertical line at *α* = 0 marks the Hopf bifurcation boundary, which governs phonation onset and offset; the dashed line marks the saddle node in a limit cycle (SNILC) boundary. Sound is produced within the oscillating regime (note symbol); silence (∅) lies outside. (c, d) Pitch (c) and normalized amplitude (d) across the phonating regime. *β* (relative) denotes the normalized distance of *β* from its minimum value to the SNILC boundary at each *α*, such that *β* (relative) = 1 corresponds to the SNILC. The white curve marks the 2200 Hz isopitch contour as an example. Pitch and amplitude vary continuously and jointly across (*α, β*) space, each depending on both biomechanical control parameters.

Pitch and amplitude vary continuously and jointly across (*α, β*) space (**Fig. 1**). Recovering *α*(t) and *β*(t) from an acoustic recording therefore yields the temporal trajectories of the biomechanical control parameters that govern song production. The inversion of the waveform to (*α, β*) is physiologically grounded, with parameter traces recovered from song correlating with simultaneously recorded air sac pressure and syringeal electromyography (*r* = 0.91 and 0.72, respectively; [11]). Subsequent automated pipelines reduced the parameter space by fixing *α* to a binary phonation gate and recovering amplitude from the rectified envelope [6, 14], making large-scale analysis tractable at the cost of discarding the continuous pressure variable. No software package yet provides automated fitting of both parameters continuously from the waveform.

We present Vocal acoustic Inversion to Biomechanical Estimates (VIBE), an open-source package that fits *α*(*t*) and *β*(*t*) against the raw acoustic waveform through iterative optimization of the normal-form equations, with integrated automated syllable segmentation. Treating *α* as a continuous free parameter rather than a phonation gate recovers a physiologically grounded air sac pressure proxy and yields better-constrained *β* estimates, as the two parameters are no longer artificially decoupled (**Fig. 1b**). We validate VIBE in twelve zebra finches with simultaneous recordings of song and air sac pressure. In an additional five zebra finches with simultaneous song and RA single-unit population recordings, we show that the VIBE-extracted parameters are encoded by RA and explain variance in RA activity that the acoustic features of song do not, consistent with the notion that RA controls vocal motor output in biomechanical rather than acoustic coordinates.

## Materials and Methods

### Biomechanical model of song production

The biomechanical model represents song production as a source-filter system, in which a nonlinear syringeal source generates sound that a passive vocal tract filter then shapes. We integrate the two as a single five-dimensional system

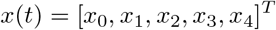

in which *x*_0_ and *x*_1_ describe the syringeal source and *x*_2_, *x*_3_, and *x*_4_ the vocal tract filter.

#### Syringeal source

The model represents each syringeal labium as a nonlinear oscillator whose midpoint displacement *x*_0_ is sustained by a surface wave propagating along the labial tissue, as in human vocal folds [7, 15]. Reducing this oscillator to its normal form around the Takens-Bogdanov bifurcation [12, 13] yields the source dynamics:

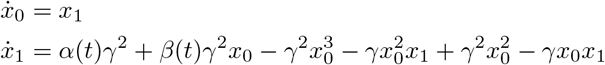

where *x*_0_ is the labial displacement, *x*_1_ is its velocity, *γ* is a fixed timescale constant (Table 1), and *α* and *β* are the two control parameters representing air sac pressure and syringeal muscle tension, respectively. The normal form retains only the terms needed to reproduce the oscillation-generating bifurcations, at low computational cost.

**Table 1.** **Model parameters**, as in Arneodo et al. [6]

| Parameter | Value | Description |
| --- | --- | --- |
| $\gamma$ | 24,000 | Oscillator timescale constant |
| $\beta_{\min}$ | $-1/3$ | Minimum syringeal muscle tension ($\beta$ lower bound) |
| $L$ | 3.5 cm | Tracheal tube length |
| $c$ | 35,000 cm/s | Speed of sound |
| $r$ | 0.1 | Tracheal reflection coefficient |
| $\kappa$ | 0.5 | Tracheal input coupling coefficient |
| $C_h^{-1}$ | $4.5 \times 10^{10}$ | OEC compliance |
| $L_b$ | $10^4$ | Beak inductance |
| $L_g$ | 82 | Glottal inductance |
| $R_b$ | $5 \times 10^6$ | Beak radiation resistance |
| $R_h$ | $6 \times 10^5$ | OEC resistance |

#### Vocal tract filter

The pressure fluctuation produced by the oscillating labia propagates through the trachea, modeled as a one-dimensional tube of length *L*, closed at the syringeal end and open at the distal end [10]. The pressure at the tracheal input combines the source contribution with the wave reflected at the open end, which returns after the round-trip delay *τ* = 2*L/c*, with speed of sound *c* [9]:

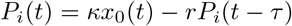

where *κ* is the input coupling coefficient and *r* is the reflection coefficient (Table 1).

To integrate the vocal tract jointly with the source in the time domain, it is represented as an equivalent electrical circuit following Perl et al. [11], the standard acoustic analog in which pressure maps to electric potential, volume flow to current, the OEC compliance to a capacitor *C*_*h*_, the glottal and beak constrictions to inductors *L*_*g*_ and *L*_*b*_, and dissipation to resistors *R*_*h*_ and *R*_*b*_. The pressure at the tracheal output relates linearly to *V*_ext_, the potential driving this circuit [10]:

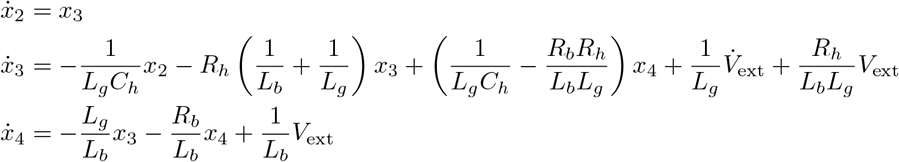

Component values are given in Table 1. The model’s acoustic output is given by the radiated pressure at the beak, *R*_*b*_ *· x*_4_(*t*).

### Algorithm overview

The algorithm extracts the time-varying control parameters *α*(*t*) and *β*(*t*) from the song waveform by decomposing each voiced frame’s fitting problem into two sequential constraints (**Fig. 2**). Observed fundamental frequency *f*_0_(*t*) at each frame constrains the parameter pair (*α, β*_rel_) to lie on a one-dimensional isopitch contour in a pre-computed parameter grid. Observed amplitude then selects the specific point on that contour. A multi-start L-BFGS-B optimization routine maximizes the Pearson correlation between the model’s predicted amplitude trajectory and the observed amplitude, subject to a smoothness regularizer on contour-position displacements between adjacent frames. VIBE (v1.0) is implemented in Python and is openly available at https://github.com/laurenostrowski/VIBE.

**Figure 2.**
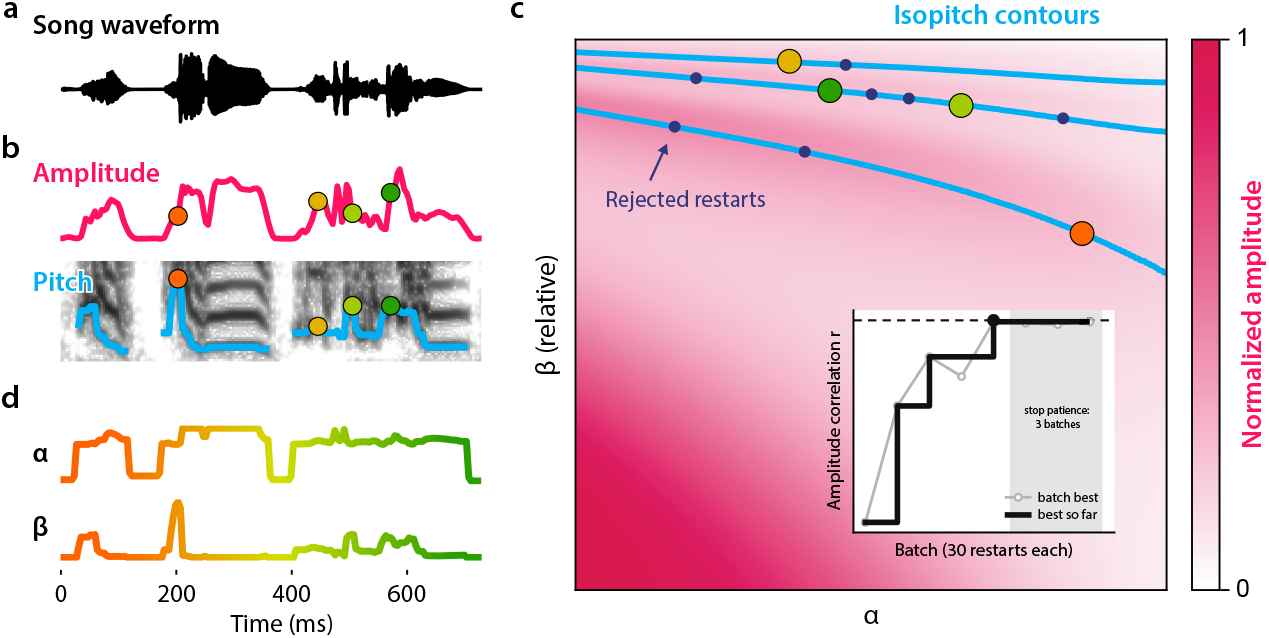
VIBE recovers biomechanical control parameters from song. (a) Song waveform excerpt spanning a few syllables. (b) Normalized amplitude envelope (top) and pitch (bottom, blue) are first estimated from the song. Colored markers indicate corresponding time points across panels. (c) For each voiced frame, the estimated pitch constrains (*α, β*) to a one-dimensional isopitch contour in precomputed parameter space (blue curves). VIBE then solves for the position along each isopitch contour that maximizes the Pearson correlation between the reconstructed and observed amplitude envelopes across all voiced frames, subject to a smoothness penalty on contour-position changes between adjacent frames. Pink shading indicates model amplitude across parameter space. Dark purple points denote (*α, β*) values from rejected optimization restarts; colored points denote the accepted solution. Inset, convergence of the multistart L-BFGS-B optimization. Each batch runs 30 restarts; the running best amplitude correlation *r* (black) and per-batch best *r* (gray) are shown across batches, with early stopping after 3 consecutive batches without improvement. (d) Recovered *α*(t) and *β*(t) trajectories across the song.

### *β* parameterization

Rather than optimizing *β* directly, the algorithm optimizes a normalized coordinate *β*_rel_ *∈* [0, 1) defined by

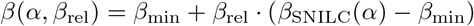

where *β*_min_ = *−*1*/*3 is the lowest physically valid *β* and *β*_SNILC_(*α*) is the SNILC bifurcation boundary, beyond which sustained oscillation is lost, whose analytic form follows from the normal-form equations [5]. *β*_rel_ thus spans *β* from its physical minimum (*β*_rel_ = 0) to the SNILC boundary, which it approaches but never reaches.

### Precomputed parameter grid

Steady-state pitch and peak-to-peak oscillation amplitude of the normal-form ODE model were precomputed over a rectangular grid spanning *α ∈* [*−*0.50, *−*0.12] (500 uniformly spaced values, restricted to the oscillating regime *α <* 0) and *β*_rel_ *∈* [0, 0.995] (500 logarithmically spaced values, denser near the SNILC boundary). At each grid point, *β* was obtained from *β*_rel_ via the normalized-coordinate mapping, the source ODE was integrated to its stable limit cycle, and the pitch and peak-to-peak amplitude of the steady-state oscillation were recorded, yielding pitch and amplitude grids over (*α, β*_rel_). The amplitude grid was normalized to [0, 1] globally.

### Algorithm application

#### Syllable detection

Song audio is segmented into syllables with the dynamic-thresholding algorithm from the AVGN pipeline [16] (https://github.com/timsainb/AVGN), which requires no manual labeling.

#### Pitch estimation

Pitch is estimated at every voiced frame using a Viterbi tracker operating on prominent peaks of the log-magnitude spectrum *M* (*k, t*) = 20 log_10_ |*S*(*k, t*) |, where *S*(*k, t*) is the complex short-time Fourier transform (STFT) coefficient at frequency bin *k* and frame *t*, computed with a Hann window and user-specified frame rate, window length, and FFT size. At each frame, candidate spectral peaks are the local maxima of *M* (*k, t*) within `[f0_min, f0_max]` returned by `scipy.signal.find` peaks, retained only if *M* (*k, t*) exceeds a per-bin noise floor (the median magnitude across silent frames) by 6 dB. Modifiable extraction parameters are the search-frequency bounds, a low-frequency bias exponent, a harmonic bonus, and the minimum peak prominence (defaults: `f0_min` = 300 Hz, `f0_max` = 4000 Hz, `freq_boost_exp = 1.0, harmonic_bonus = 5 dB, min_prominence dB = 5 dB)`. A visualization routine overlays the estimated pitch contour on the spectrogram so the user can verify and adjust these parameters before committing them to the full pipeline; parameters tuned this way should yield consistent pitch estimates across a given bird’s entire song corpus.

Viterbi decoding selects the sequence of candidate peaks maximizing the total path score, comprising a per-frame emission score with three additive terms in dB,

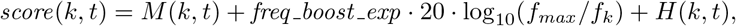

where *f*_*k*_ is the frequency of bin *k*, the second term is a log-scale bias favoring lower fundamental frequencies (`freq_boost_exp` = 1.0 by default), and *H*(*k, t*) adds 5 dB per harmonic (2*f*_*k*_ through 5*f*_*k*_) present as a peak within ± 15% frequency tolerance in the same frame. The transition cost between consecutive voiced frames is *−* 0.01*·* |Δ*f*|, penalizing frequency jumps. The optimal path is recovered by backtracking from the highest-scoring terminal state. The trajectory is then smoothed by segmenting at inter-frame jumps exceeding 500 Hz and applying a 5-sample median filter independently within each segment of at least five frames. Residual gaps at voiced frames are filled by averaging the nearest valid neighbors on each side (or copying the single available neighbor); voiced frames with no valid neighbor in either direction are reclassified as unvoiced.

#### Amplitude estimation

Per-frame amplitude is computed as the square root of total STFT power in the 200–8000 Hz frequency band,

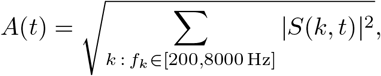

where |*S*(*k, t*) |^2^ is the power at bin *k* and frame *t*. The resulting amplitude trace is normalized per song. In most experimental settings the microphone is not fixed to the bird, so absolute amplitude across songs is confounded by the bird’s distance and orientation relative to the microphone. Within-song amplitude fluctuations, by contrast, mainly reflect vocal output, as birds typically perch in a single position while singing.

#### Isopitch contour extraction

For each voiced frame *t* with estimated pitch *f*_0_(*t*), the isopitch contour *C*_*t*_ (**Fig. 1c–d**) is extracted from the precomputed pitch grid as the locus of (*α, β*_rel_) pairs at which the grid pitch equals *f*_0_(*t*), via marching squares (`matplotlib` contour). Each contour is an ordered sequence of vertices parameterized by normalized vertex index *p* ∈ [0, 1]. Model amplitude along *C*_*t*_ is obtained by bilinear interpolation of the amplitude grid at each vertex, with linear extrapolation beyond the grid bounds. Contour extraction and amplitude interpolation are parallelized across voiced frames.

#### Objective function

The optimization variable is the vector of per-frame contour positions *p* = {*p*_*t*_ : *t* ∈ voiced}, with *p*_*t*_ ∈ [0, 1]. For a given *p*, the model amplitude *Â*(*t*) at each voiced frame is obtained by linear interpolation along *C*_*t*_ at position *p*_*t*_. The objective is

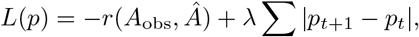

where *r* is the Pearson correlation between observed amplitude *A*_obs_ and model amplitude *Â* over all voiced frames, and the smoothness term sums absolute position differences over temporally adjacent voiced frames (*t* and *t* + 1 both voiced and consecutive in STFT frame index, so inter-syllable silences do not contribute), with weight *λ* = 0.001. Pearson correlation makes the objective invariant to the absolute scale and offset of the amplitude, so the optimization fits the shape of the amplitude trajectory rather than its absolute value.

#### Initialization

Isopitch contour positions are initialized by a rank-based mapping that gives consistent initialization across songs: the percentile rank of *A*_obs_(*t*) among all voiced-frame amplitudes is computed, and each *p*_*t*_ is set to the contour vertex whose model amplitude holds that same rank along *C*_*t*_. This initialization approximates the observed amplitude profile, so the optimizer begins close to an amplitude-consistent solution.

#### Multi-start L-BFGS-B optimization

*L*(*p*) is minimized by box-constrained quasi-Newton optimization (L-BFGS-B; [17]) with *p*_*t*_ ∈ [0, 1], maximum 1000 iterations, and convergence tolerance `ftol` = 10^*−*8^ per trial. To manage non-convexity, optimization proceeds in batches of 30 independent trials. Apart from the first trial, which uses the rank-based initialization above, each trial draws *p*_*t*_ independently from a uniform distribution over [0.2, 0.8], smoothed with a 5-sample moving average to reduce initialization discontinuities. The lowest-*L* result in each batch is retained and compared to the running global best. Batching continues until either 10 batches complete or 3 consecutive batches fail to improve the global best by more than 10^*−*3^. The best fit solution gives *α*(*t*) and *β*(*t*) at every voiced frame.

#### Silent frames

At silent frames, *β*_rel_ is fixed at 0.99 and *α* decays with the duration of the silence as *α*(Δ) = 0.5 + 1.05*·* exp(*−* Δ*/τ* ), with Δ the silence duration and *τ* = 0.030 s. These coefficients are fit to air sac pressure recorded during the silent inspirations between syllables (minibreaths; [18]). Phonation boundaries are smoothed with a 5 ms raised-cosine ramp.

### Song synthesis

Synthetic waveforms are generated by forward integration of the coupled syringeal source and vocal tract ODEs. The fitted control parameters *α*(*t*) and *β*(*t*) are upsampled from frame rate to audio rate, and the model is integrated forward with fourth-order Runge-Kutta, oversampled eightfold. Synthesis is parallelized across windows, on CPU by multiprocessing or on GPU by batching.

### Air sac pressure dataset

Air sac pressure recordings were collected from twelve zebra finches, in accordance with a protocol approved and supervised by the Institutional Animal Care and Use Committees at Texas Christian University and the University of Utah. The dataset comprises 44 songs with paired air sac pressure recordings across these birds: 35 songs from three birds and one each from the remaining nine.

#### Animal husbandry

All finches were maintained on a 14-10h light cycle. Before experimental testing, birds were housed in communal cages with 4-12 birds per cage. During experiments, the birds were individually housed in small cages (31.8× 10.5 ×25.4 cm) contained in a sound-attenuating box (78.7× 33 ×33 cm). Four sides of the sound-attenuating box were lined with 1-in. thick acoustic foam (Auralex Acoustics) to dampen acoustic reflections, and the front of the box was open for conspecific presentation and acoustic interaction to promote singing. A microphone was suspended 14 cm above or in front of the center of the perch.

#### Surgical procedure

Air pressure was recorded via a small cannula (Silastic tubing, 0.76 mm I.D., Dow Corning) inserted into the left or right cranial thoracic air sac (**Fig. 3**). All procedures were performed under aseptic surgical conditions and general anesthesia (1-2% isoflurane in oxygen). A small opening was made in the body wall below the last rib; the cannula was sutured to the body wall and sealed to the skin with tissue adhesive. Its free end was connected to a pressure transducer (Fujikura XFHM-02PGR). A 50/50 mixture of lidocaine (2%) and bupivacaine (0.5%) was applied topically as a postsurgical analgesic. The transducer was centered between the wings and held in place with a Velcro tab attached to an elastic band. The transducer weight was offset by a counterweighted balance arm that allowed free movement in the cage. Birds were monitored postoperatively until they could perch, hop, and move freely while carrying the transducer and wires.

**Figure 3.**
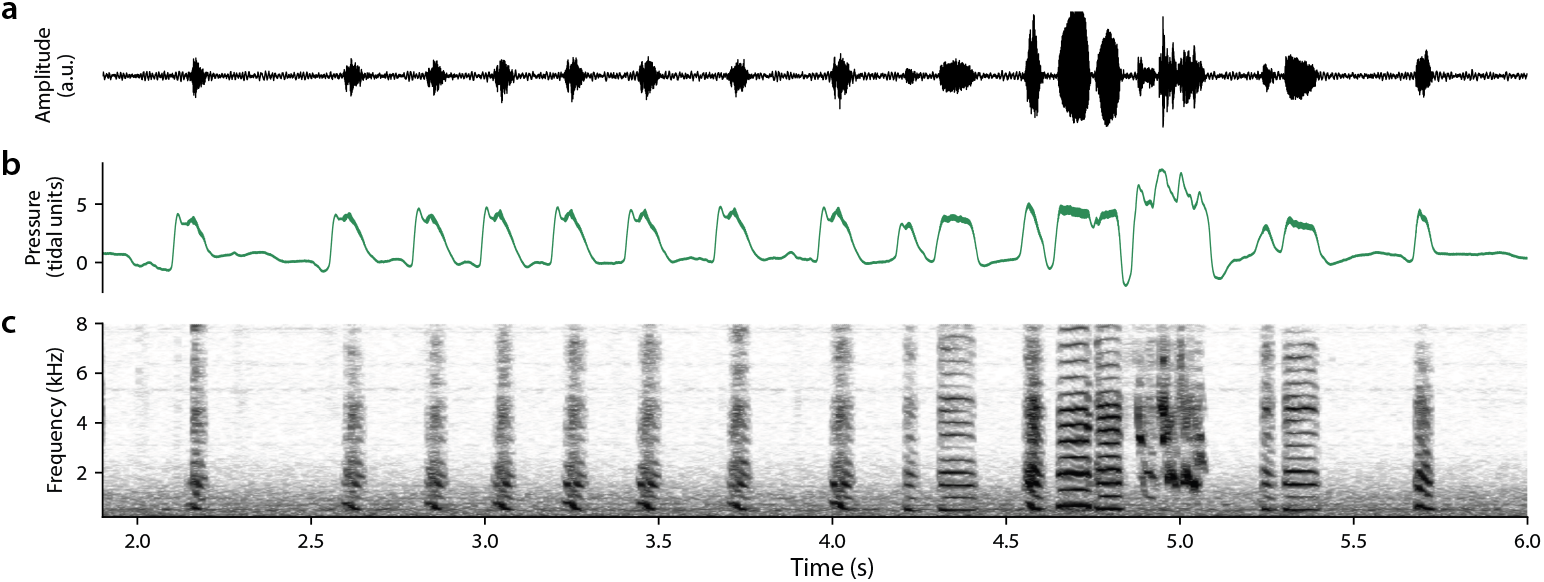
Simultaneous song and air sac pressure recording. (a) Song waveform, (b) subsyringeal air sac pressure, and (c) song spectrogram, recorded simultaneously. Pressure is normalized to tidal volume, such that 0 corresponds to ambient pressure during quiet respiration; expiratory pulses driving each vocalization appear as positive deflections.

#### Data collection

Signals from the backpack-mounted pressure transducer were passed to a DC amplifier (gain 100, 6 kHz low-pass; Brownlee Precision Model 440) and recorded together with song. A microphone (Earthworks TC20 or SR20) was positioned 14 cm in front of or above the bird; its signal was amplified and high-pass filtered (300 Hz) before digitization. Song and air pressure were digitized with a National Instruments NI USB-6251 board and Avisoft Recorder software (Avisoft Bioacoustics), and stored as multichannel 16-bit WAV files at 22.05 or 44.1 kHz. Acquisition was triggered whenever air pressure exceeded a user-defined threshold set at approximately twice the peak of quiet respiration. Each file spanned 5 s before threshold onset to 5 s after it was last exceeded, giving at least 10 s of paired respiration and song.

### RA dataset

Electrophysiological neural recordings were collected from five zebra finches, in accordance with a protocol approved and supervised by the Institutional Animal Care and Use Committee at the University of California San Diego. The dataset comprises 398 songs with paired neural recordings across these birds, with a mean of 80 songs (range 35-131) and 184 neurons (range 88-280) per bird. Animal husbandry, surgery, electrophysiological recording, and histology followed procedures described in detail previously [19], summarized below.

#### Animal husbandry

Prior to surgery, zebra finches were acclimated to commercial acoustically isolated chambers (Eckel) on a 14-10h light cycle. Each chamber was fitted with a UMA-8 V2.0 multichannel USB microphone array (miniDSP) for continuous audio capture and acoustic panels for sound attenuation. Audio was monitored continuously during daylight hours and mirrored hourly to a shared server, where custom Python signal-processing routines parsed the recordings and counted each bird’s daily song renditions. Birds producing the most songs per day were selected for neural implantation.

#### Surgical procedure

Zebra finches were implanted with a multi-shank Neuropixels 2.0 probe (Imec) targeting RA (**Fig. 4c**). All procedures were performed under aseptic conditions and general anesthesia (1–2% isoflurane in oxygen). The day before surgery, birds received intramuscular enrofloxacin (Enrosite; 20 mg/kg) and dexamethasone (Dexasone; 5 mg/kg); intramuscular carprofen (Rimadyl; 4 mg/kg) was administered postoperatively as needed for pain and inflammation. Birds were head-fixed in a digital stereotaxic frame, the head feathers plucked, and the scalp disinfected with betadine. A midline incision from the base of the beak to the back of the head exposed the cranium. The head was angled at 130°, defined as the angle between the horizon and the line from the beak to the ear. A craniotomy and durotomy centered 2200 µm lateral and 300 µm anterior to the Y-sinus exposed the brain. A Neuropixels probe, held by a custom shaft and stained with DiI (DiIC18(3)) for post-mortem track reconstruction, was inserted to a depth of 3500 µm. Recordings from 384 sites at 15 µm pitch along the probe spanned the full dorsoventral extent of RA [3, 20]. The probe- and headstage-holding shafts were fixed to the skull with adhesive cement (C&B Metabond), the durotomy was covered with silicone dural substitute (Dow Corning 3-4680), and the skin was sutured around the implant. Total implant weight was 1.2 ± 0.2 g. Implantation at RA was confirmed post-mortem by histological assessment of the probe track. All five birds were implanted in the right hemisphere.

**Figure 4.**
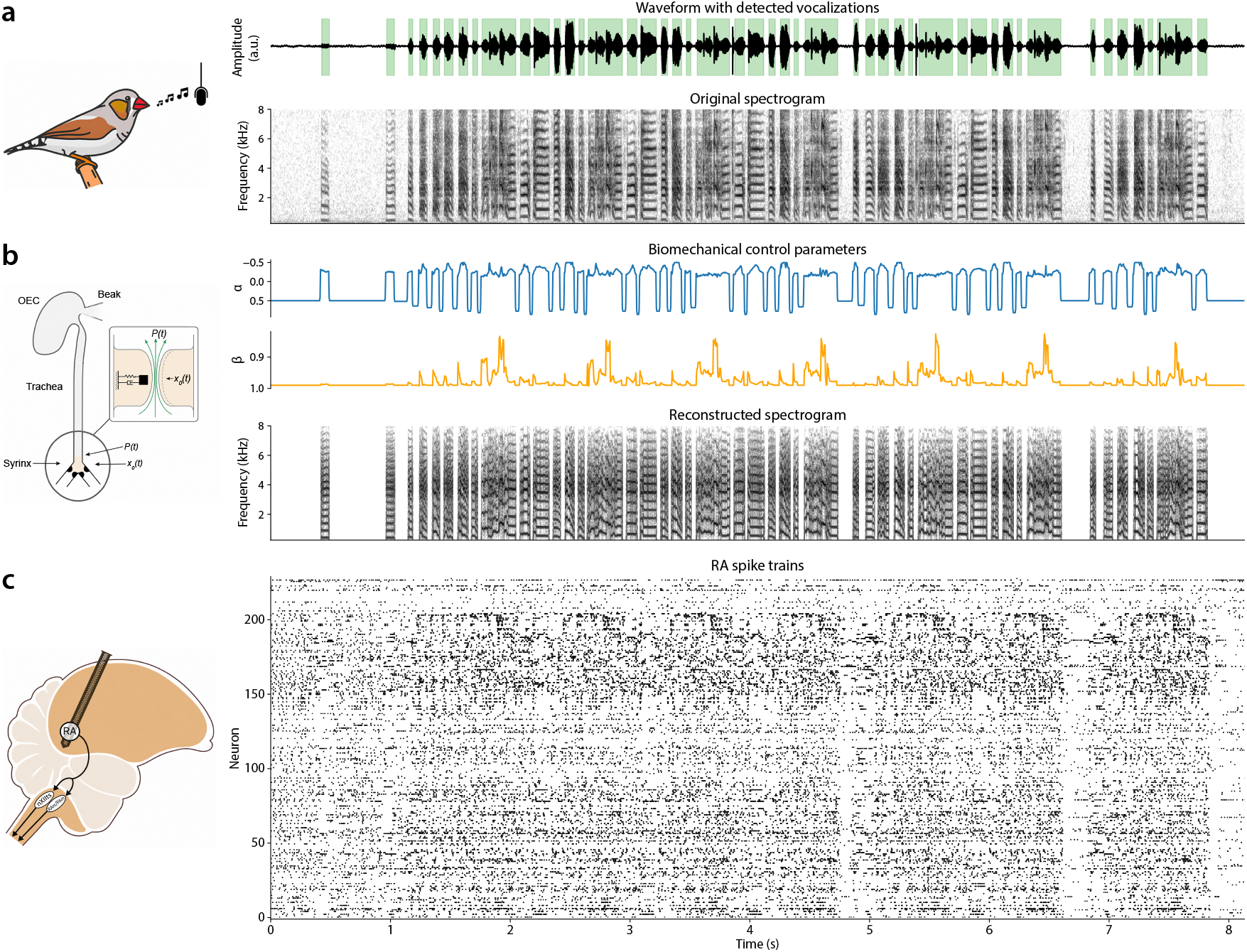
Song, recovered biomechanical control parameters, and simultaneous RA population activity. (a) Song waveform (top; amplitude in arbitrary units) with detected vocalizations shaded in green, and the corresponding spectrogram (bottom). (b) Left, schematic of the vocal organ: the syringeal labia (*x*_0_(*t*)) are driven by subsyringeal pressure *P* (*t*), and source sound passes through the trachea and oropharyngeal-esophageal cavity (OEC) before radiating from the beak. Right, biomechanical control parameters *α*(*t*) (air sac pressure, blue) and *β*(*t*) (syringeal muscle tension, orange) recovered from the waveform by VIBE, and the spectrogram reconstructed from these parameters (bottom). (c) Left, sagittal schematic showing the Neuropixels probe targeting RA. Right, spike trains of n = 232 simultaneously recorded RA neurons during the song in (a).

#### Data collection

Recording of simultaneous neural and song data began the morning after implantation. Implant weight was offset by a 1.2 g counterweight routed over a pulley on thin nylon wire. Each session comprised three continuous hours of recording, with a female conspecific introduced for the final hour, to elicit both undirected and directed song in the same session. Voltage signals from 384 Neuropixels channels were amplified, multiplexed, and digitized on the probe base, then relayed via the headstage to a Neuropixels PXIe acquisition module (Imec) in a PXIe-1083 chassis (National Instruments) and recorded with SpikeGLX (v.20240129, Imec phase30 API v3.62.1; https://billkarsh.github.io/SpikeGLX/). Continuous full-band voltage waveforms from all channels were saved at 30 kHz. Audio was acquired with a TC20 omnidirectional microphone (Earthworks) at 40 kHz using a PXIe-6363 multifunction data acquisition module (National Instruments). Neural and audio signals were synchronized by a common 1 Hz clock from the Neuropixels PXIe acquisition module.

#### Histology

Following physiological recording, birds were transcardially perfused with heparinized saline followed by 4% paraformaldehyde (PFA) in phosphate buffer. Brains were then extracted and post-fixed in 4% PFA for at least 24 hours at 4°C. For cryoprotection, they were transferred to 30% w/v sucrose 72 hours before sectioning. The specimen was embedded in Tissue-Tek O.C.T. compound on the cryostat stage. 30 µm sagittal sections were cut and mounted on glass slides. Sections were examined by light microscopy and imaged with a digital camera.

#### Neural signals processing

All data processing and analysis were performed in Python 3 using custom software unless otherwise noted. Putative action potentials (hereafter, spikes) were extracted from continuous extracellular waveforms and sorted into putative single neurons (units) using Kilosort4 via SpikeInterface with default preprocessing parameters. Sorted units were pre-screened using three metrics computed in SpikeInterface, the interspike-interval (ISI) violation ratio, signal-to-noise ratio, and presence ratio. Every unit was then manually reviewed and classified as single-unit activity (SUA), multi-unit activity (MUA), or noise on the basis of waveform shape, autocorrelogram, and amplitude stability across the session. Across the 5 birds, this yielded 918 SUA, 242 MUA, and 304 noise clusters (mean 184 SUA per bird, range 88-280). Only single units were retained for subsequent analysis.

#### Audio processing

Songs were automatically detected in each recording session using custom song-recognition software. VIBE was then used to fit the biomechanical model to each song, recovering *α*(*t*) and *β*(*t*) (**Fig. 4b**).

### Data analysis

#### End-to-end reconstruction validation

To validate VIBE, we resynthesized each song by integrating the recovered *α*(*t*) and *β*(*t*) through the source-filter model, and compared the reconstructed song to the original. We quantified reconstruction quality by pitch correlation, amplitude correlation, and spectrogram earth mover’s distance (EMD).

Pitch and amplitude correlations were computed across frames with vocalizations as the Pearson correlation between the reconstructed and original per-frame pitch and amplitude envelopes, respectively. Amplitude was the square root of STFT power integrated over 200–8000 Hz. Pitch correlation is unity by construction, as the algorithm constrains each frame to an isopitch contour.

To assess spectral reconstruction beyond pitch and amplitude, we computed the earth mover’s distance [21], as used previously to compare synthetic and recorded birdsong spectrograms [6]. Magnitude spectrograms were computed with a 16 ms Hann window, 4 ms hop, and a 1024-point FFT. Each frame was normalized to unit mass, and the per-frame EMD was computed as the one-dimensional Wasserstein distance between the reconstructed and original spectra over frequency, making it robust to small pitch offsets that would otherwise inflate bin-wise errors, then averaged across frames. To verify that the reconstruction captured the temporal structure of each song, we compared the observed EMD to a cyclic-shift null. To compute a per-song null, the reconstruction was cyclically shifted by 200 random offsets (minimum 10% of song length), the EMD recomputed at each, and the null taken as the median across shifts. Specificity was quantified as ΔEMD = EMD_null_ *−* EMD_observed_, with positive values indicating lower EMD than time-shifted controls, and compared to zero for each bird (Wilcoxon signed-rank test).

For visualization (**Fig. 5a**), we additionally reconstructed each song from a random starting configuration, in which each frame was assigned a uniformly random position along its isopitch contour, and from the rank-based initialization prior to optimization.

**Figure 5.**
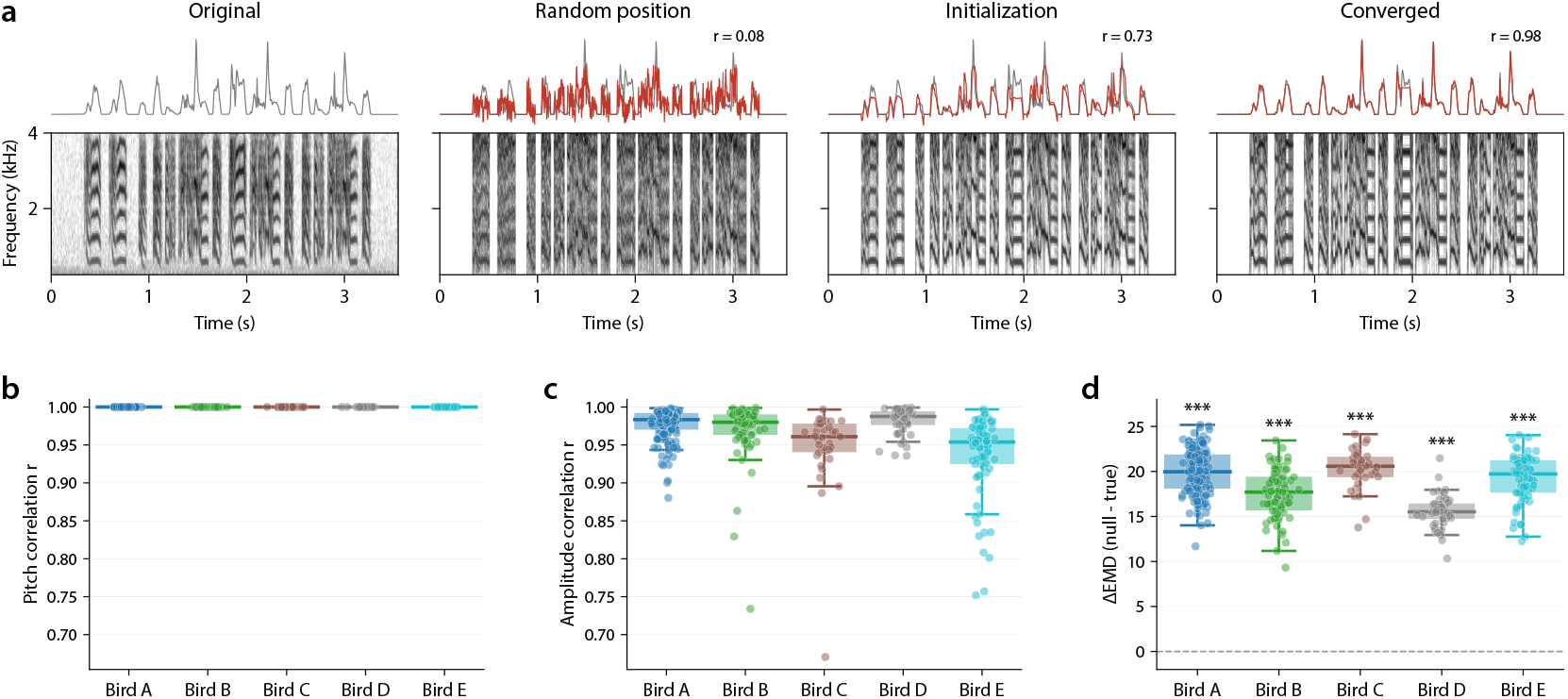
End-to-end reconstruction validates the recovered parameters. VIBE recovers parameters by deterministic inversion of each song, so reconstruction accuracy is itself the validation. (a) Reconstruction of a representative song (the song with the median amplitude-correlation score across Bird B’s songs). Columns show the original, a random position on the isopitch contours (shown for comparison; not used in fitting), the rank-based initialization, and the converged fit. Top, amplitude envelope of the original (gray) and reconstruction (red), with Pearson correlation (*r*) annotated; bottom, corresponding spectrograms. (b) Pitch correlation between reconstructed and original song, per song, grouped by bird. Correlation is exactly unity by construction: the algorithm constrains each frame to an isopitch contour, so recovered pitch equals observed pitch. (c) Amplitude correlation between reconstructed and original amplitude envelopes, per song, grouped by bird. (d) Reconstruction specificity, quantified as the difference in earth mover’s distance (ΔEMD) between a cyclic-shift null and the true reconstruction (null *−* true; positive values indicate closer agreement with the original than shifted controls). For each song, the reconstructed spectrogram was cyclic-shifted relative to the original (200 shifts, minimum 10% of song length) and the per-frame EMD recomputed; the null is the median across shifts. ΔEMD was significantly positive in all birds (Wilcoxon signed-rank test). Boxes show median and interquartile range across songs; whiskers extend to the most extreme points within 1.5 times the interquartile range. \*\*\**p <* 0.001

### Air sac pressure analysis

We fit *α*(*t*) and *β*(*t*) to every song waveform using VIBE and assessed the correspondence between the recovered *α* and simultaneously recorded air sac pressure. The song amplitude envelope served as a benchmark, allowing us to test whether *α* predicts air sac pressure beyond what this readily available acoustic feature captures. Amplitude was computed as the square root of STFT power integrated over 200–8000 Hz. All comparisons were conducted over both the full song and frames with vocalizations only.

We applied four tests, each defined so that a more positive value indicates that *α* better explains recorded air sac pressure. First, to assess whether *α* predicts pressure above chance, we used a cyclic-shift permutation test (1000 shifts) with statistic median (MSE_null_ *−* MSE_*α*_). Second, to assess whether *α* predicts pressure more accurately than amplitude, we computed (MSE_amplitude_ *−* MSE_*α*_), where each term is the MSE of a per-song linear fit. Third, to assess whether *α* accounts for variance in pressure not explained by amplitude, we compared two nested linear models, amplitude alone and amplitude and *α* jointly, using (MSE_amplitude_ *−* MSE_amplitude, *α*_) evaluated on held-out frames. Fourth, to assess whether *α* predicts pressure more accurately than a binary phonation gate, we computed (MSE_gate_ *−* MSE_*α*_), where the gate is a two-level indicator of phonation versus silence fit to pressure per song. This gate corresponds to the reduced parameterization in which *α* is fixed to a binary phonation state, as used in prior automated pipelines [6, 14].

Because the gate is constant within each vocalization, this comparison was evaluated on the within-vocalization pressure residual, obtained by subtracting each vocalization’s mean pressure. To account for the non-independence of songs within birds, we modeled each per-song statistic with a linear mixed-effects model (fixed intercept; random intercept for bird) and tested whether the grand mean differed from zero, reporting the fixed-effect estimate and 95% confidence interval. Per-song values for the binary gate comparison were right-skewed, so its intercept and confidence interval were estimated on signed-log-transformed values to meet the model’s normality assumption, then back-transformed to the original units. As a descriptive measure, we report the Pearson correlation between *α* and recorded pressure across songs (mean *±* SD across birds).

### RA analysis

To determine whether RA activity encodes the recovered biomechanical control parameters, and whether it does so beyond the acoustic features they produce, we predicted single-unit spiking from *α, β*, amplitude, and pitch. For each song, the biomechanical control parameters (*α, β*) and acoustic features (amplitude envelope, pitch) were represented at 4 ms resolution, and spike trains were binned to match. Data were concatenated across songs, with time-shifts and cross-validation folds constrained to respect song boundaries. For each neuron, we fit Poisson generalized linear models (GLMs) predicting binned spike counts from the vocal parameters. Performance was evaluated by 5-fold cross-validation with folds assigned at the song level, such that temporally correlated frames within a song were confined to a single fold. Held-out log-likelihood was summed across test folds.

To determine whether individual neurons encode the biomechanical control parameters, we fit three Poisson GLMs per neuron: *α* alone, *β* alone, and *α* and *β* jointly. Significance was assessed against a cyclic-shift null. For each of 200 shifts, predictors within each song were time-shifted by a random offset (shared across *α* and *β* in the joint model), the model refit, and the held-out log-likelihood recomputed. One-sided p-values were computed as the fraction of null log-likelihoods exceeding the observed value.

P-values were corrected across neurons within each bird using the Benjamini-Hochberg procedure at FDR *<* 0.05. The fraction of neurons significantly encoding each parameter is reported per bird.

To test whether *α* and *β* carry information about RA activity beyond what acoustic features capture, we compared nested Poisson GLMs. For each neuron and each acoustic reference (amplitude, pitch, or both), we first identified responsive neurons as those whose spike counts were significantly predicted by the acoustic feature alone, using the same cyclic-shift null procedure as above (200 shifts; Benjamini-Hochberg FDR *<* 0.05 across neurons within each bird). In responsive neurons, *α* and *β* were added to the acoustic (reduced) model, and the improvement of the full model was quantified by the change in Akaike information criterion computed from held-out log-likelihoods, ΔAIC = 2*k −* 2(LL_full_ *−* LL_reduced_), with *k* = 2 added parameters; negative ΔAIC indicates that adding *α* and *β* improved held-out prediction. Significance was assessed against a cyclic-shift null in which *α* and *β* were jointly shifted within songs. For each of 200 shifts, ΔAIC was recomputed, and the one-sided p-value was computed as the proportion of null ΔAIC values below the observed value. The fraction of responsive neurons for which *α* and *β* significantly improved prediction is reported per bird.

## Results

### End-to-end reconstruction accuracy

To assess how faithfully the recovered parameters capture the original song, we resynthesized each song from its fitted *α* and *β* and compared the reconstruction to the original, in the five RA-dataset birds with sufficient songs for reliable per-bird estimates, along three axes: pitch, amplitude, and spectral content (**Fig. 5a**). Because VIBE recovers parameters by deterministic inversion, the accuracy with which the recovered *α*(t) and *β*(t) reconstruct the original song is used as validation.

Reconstruction quality improved over optimization, and the converged fit closely matched the original. Pitch correlation was unity by construction, as each frame is constrained to an isopitch contour (**Fig. 5b**). Amplitude was recovered with high fidelity across birds (median Pearson *r* = 0.95 to 0.99 across the five birds; **Fig. 5c**). To assess spectral reconstruction beyond pitch and amplitude, we computed the earth mover’s distance between reconstructed and original spectrograms and compared it to a cyclic-shift null that preserves each song’s overall spectral content but breaks its temporal alignment. Reconstructions had significantly lower EMD than the shifted null in every bird (ΔEMD significantly positive; Wilcoxon signed-rank test, *p <* 0.001 for all birds; **Fig. 5d**), confirming that VIBE reconstructs the specific spectrotemporal content of each song.

While accurate reconstruction confirms that *α* and *β* capture the acoustics of song, the definitive test is against empirical ground truth. We therefore recorded air sac pressure during singing and compared it to the recovered *α*.

### *α* provides a proxy for physiological air sac pressure

To validate that *α* recovered by VIBE reflects the physiological variable it encodes, we compared *α* to simultaneously recorded air sac pressure across 44 songs from 12 birds. Across songs, *α* closely matched the pressure trace (**Fig. 6a**), with a mean Pearson correlation of 0.87 ± 0.03 (range 0.78–0.93). To quantify this correspondence, we applied four statistical tests. The first assessed whether *α* predicts pressure above chance, two compared *α* to the acoustic amplitude envelope as a readily available benchmark, and the final compared *α* to a binary phonation gate, the parameterization used in prior automated pipelines [6, 14].

**Figure 6.**
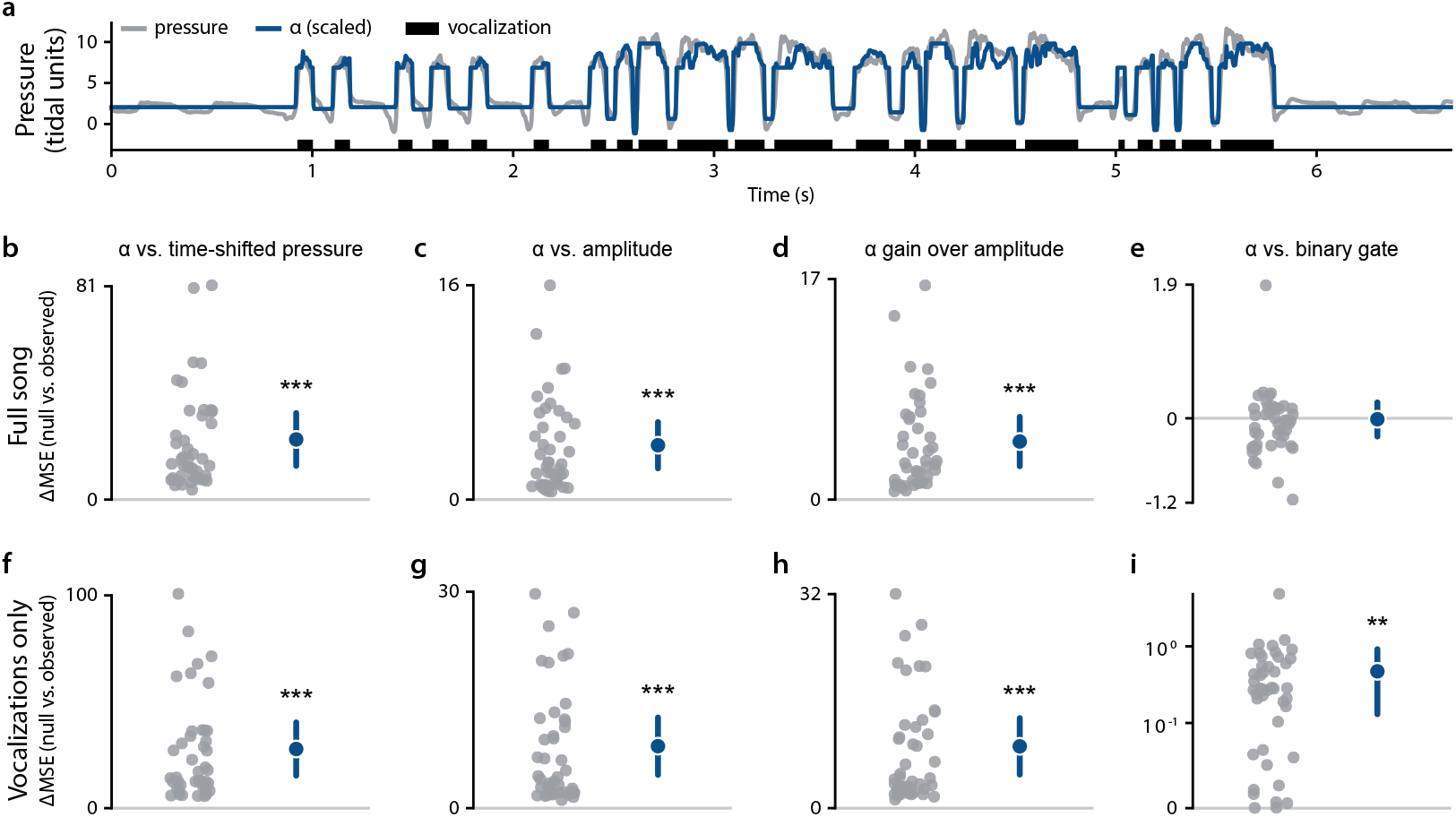
*α* recovered by VIBE is a proxy for physiological air sac pressure. (a) Recorded air sac pressure (gray) and the fitted *α* trajectory (blue), sign-aligned and scaled to pressure by least squares, for an example song (Pearson *r* = 0.93). Black bars mark vocalizations. *α* tracks the pressure trace across vocalizations and inter-vocalization silences. (b-i) Four model-comparison tests, each evaluated over full songs (b-e) and frames with vocalizations only (f-i), across 44 songs from 12 birds. Gray points are individual songs; blue points and bars show the linear mixed-effects model intercept and 95% confidence interval (bird as a random effect). Quantities in (b-d, f-h) are differences in mean squared error (ΔMSE) between predicted and recorded pressure, in squared normalized pressure units; values above zero indicate that *α* improves the prediction. (b, f) *α* predicts pressure better than null time-shifted pressure (intercept = 22.8 [12.8, 32.9] full; 27.8 [15.3, 40.3] vocalizations). (c, g) *α* predicts pressure better than acoustic amplitude (4.1 [2.3, 5.8] full; 8.6 [4.6, 12.6] vocalizations). (d, h) Adding *α* as a predictor to an amplitude-only model further reduces prediction error (4.5 [2.6, 6.4] full; 9.2 [5.0, 13.5] vocalizations). (e, i) *α* versus a binary phonation gate. As the gate is constant within each phonation, *α* was compared to it on the within-phonation pressure residual. Per-song values for the binary gate comparison (i) were right-skewed, so the intercept and 95% CI were estimated from a mixed-effects model (bird as a random effect) on signed-log-transformed values to satisfy normality, then back-transformed to ΔMSE units; i is plotted on a logarithmic y-axis, all other panels on a linear axis. *α* modestly outperformed the binary gate within vocalizations (i; 0.5 [0.1, 0.9]), with the two statistically indistinguishable across the full song (e; 0.0 [*−* 0.3, +0.2]). \*\**p <* 0.01, \*\*\**p <* 0.001; comparison without an asterisk is not significant.

*α* predicted pressure significantly better than a cyclically time-shifted null (ΔMSE = 22.8 [12.8, 32.9] squared normalized pressure units; linear mixed-effects model, *p <* 0.001; **Fig. 6b**), more accurately than amplitude alone (ΔMSE = 4.1 [2.3, 5.8]; *p <* 0.001; **Fig. 6c**), and explained significant additional variance in pressure not accounted for by amplitude when added as a second predictor (ΔMSE = 4.5 [2.6, 6.4]; *p <* 0.001; **Fig. 6d**). All three effects were larger when restricted to frames with vocalizations (ΔMSE = 27.8 [15.3, 40.3], 8.6 [4.6, 12.6], and 9.2 [5.0, 13.5], respectively; *p <* 0.001 for all; **Fig. 6f-h**), consistent with *α* being most tightly constrained by the model during phonation. *α* modestly outperformed the binary phonation gate within vocalizations (ΔMSE = 0.5 [0.1, 0.9]; *p* = 0.004; **Fig. 6i**), while the two were statistically indistinguishable across the full song (ΔMSE = 0.0 [*−* 0.3, +0.2]; *p* = 0.9; **Fig. 6e**), indicating that the advantage of continuous *α* over a binary gate lies in the graded pressure variation within a vocalization and the differences in mean pressure across vocalizations.

### RA activity encodes biomechanical control parameters

To determine whether RA activity encodes the biomechanical control parameters recovered by VIBE, we tested whether *α* and *β* predict RA activity, and whether they explain variance in RA activity beyond what the acoustic features they produce already capture. We fit Poisson GLMs to each neuron’s binned spike counts, assessing significance against a cyclic-shift null and quantifying the improvement beyond acoustic features by the change in Akaike information criterion (ΔAIC) on held-out data (Methods).

The biomechanical control parameters accounted for RA activity in a majority of neurons. *α* significantly predicted spike activity in 72% of neurons pooled across birds (Birds A-E: 91%, 44%, 58%, 91%, and 95%), *β* in 68% of neurons (87%, 44%, 41%, 88%, and 93%), and *α* and *β* jointly in 76% of neurons (94%, 51%, 55%, 95%, and 95%; **Fig. 7a**).

**Figure 7.**
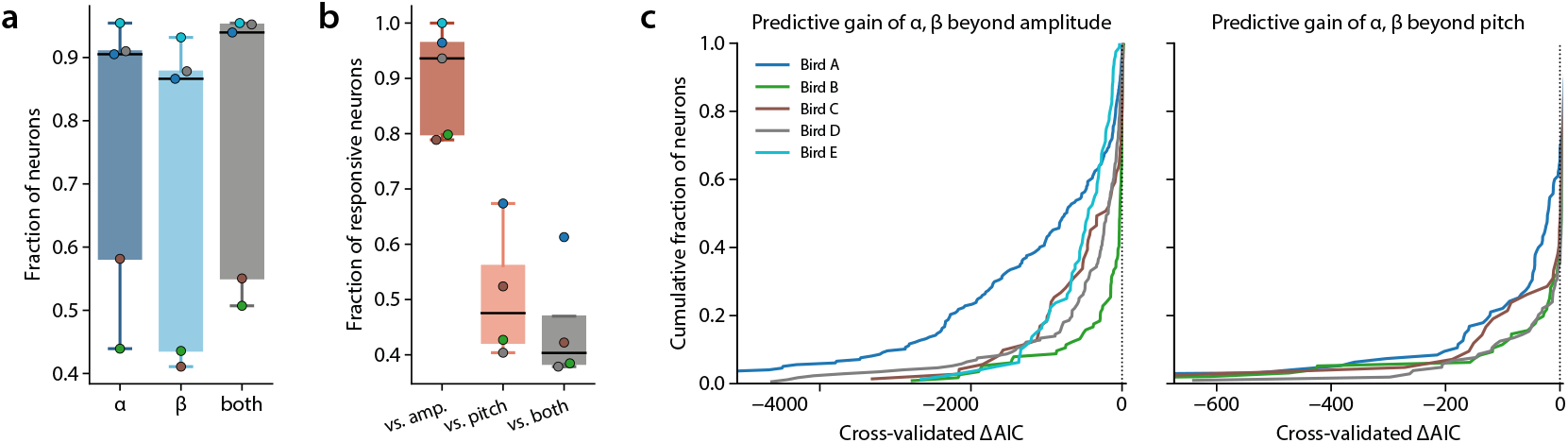
Biomechanical parameters predict RA activity beyond acoustic features. (a) Fraction of RA neurons whose spike activity was significantly predicted by *α, β*, and *α* and *β* jointly. For each neuron, a Poisson generalized linear model predicting spike counts from each parameter was compared against a cyclic-shift null distribution (200 shifts), using held-out log-likelihood as the test statistic; p-values were FDR-corrected across neurons within each bird. Each point is one bird; boxes show median and interquartile range across the five birds, with whiskers extending to the most extreme values within 1.5 times the interquartile range. (b) Fraction of responsive neurons for which adding *α* and *β* significantly improved prediction beyond amplitude, pitch, or both. Responsive neurons were those significantly predicted by the acoustic feature (cyclic-shift null, FDR-corrected across neurons); in these, *α* and *β* were added to the reduced model and the ΔAIC assessed against a cyclic-shift permutation null (200 shifts, *p <* 0.05). Plotted as in (a); Bird E, with no responsive neurons for pitch or amplitude & pitch, is omitted for those comparisons. (c) Cumulative distributions of the per-neuron cross-validated ΔAIC between the reduced model (amplitude, left; pitch, right) and the full model including *α* and *β*, one curve per bird. ΔAIC was computed from held-out log-likelihoods; negative values indicate improved prediction with *α* and *β*. Curves are truncated at the 1st percentile of the pooled distribution; 7 neurons (Bird A) fall beyond the left axis limit for amplitude, and 4 (2 from Bird A, 1 each from Birds B and C) for pitch.

We next asked whether *α* and *β* explain variance in RA activity beyond that explained by the acoustic features they produce, testing in each acoustically responsive neuron whether adding *α* and *β* to the acoustic model significantly improved held-out prediction. *α* and *β* improved prediction beyond amplitude in 91% of amplitude-responsive neurons pooled across birds (Birds A-E: 96%, 80%, 79%, 94%, and 100%; median ΔAIC = *−* 315 among improved neurons; **Fig. 7b, c, left**). *α* and *β* also improved prediction beyond pitch in 50% of pitch-responsive neurons (Birds A-D: 67%, 43%, 52%, and 40%; median ΔAIC = *−* 40; **Fig. 7b, c, right**), and beyond amplitude and pitch jointly in 45% of jointly responsive neurons (61%, 38%, 42%, and 38%; **Fig. 7b**). Bird E had no neurons responsive to pitch and is omitted from the pitch and joint comparisons. We therefore find that the biomechanical control parameters add explanatory power for RA activity beyond the acoustic features they produce, with the largest improvement over the amplitude model.

## Discussion

We present VIBE, a novel, fully automated pipeline to recover *α* and *β* as continuous free parameters from the raw song waveform. The waveform-to-parameters inversion problem is non-trivial: *α* and *β* are biomechanically coupled, and the mapping from parameter space to acoustics is nonlinear, making joint recovery substantially more challenging than fitting either parameter in isolation. Treating both as continuous free parameters with joint optimization against the song waveform preserves the coupling structure of the underlying model, recovers a physiologically grounded air sac pressure proxy, and yields pressure-constrained syringeal muscle tension estimates across the full dynamic range of song.

Validation against recorded air sac pressure across 44 songs from 12 birds confirmed that VIBE’s inversion is physiologically grounded. Recovered *α* predicted pressure above chance, outperformed the acoustic amplitude envelope, and explained variance in air sac pressure not captured by amplitude alone, and, within vocalizations, exceeded a binary phonation gate. A limitation of our dataset is the absence of syringeal muscle electromyography, precluding direct validation of *β*.

Applying VIBE to Neuropixels recordings from RA, we found that most neurons encoded the recovered parameters, and that *α* and *β* added explanatory power for RA activity beyond the acoustic features of song. The predictive gain was largest beyond the amplitude envelope, where *α* and *β* improved prediction in nearly all responsive neurons, and more modest beyond pitch. This asymmetry is consistent with the biomechanics: *α*, a proxy for air sac pressure, captures respiratory drive that the amplitude envelope indirectly reflects, whereas pitch is more directly tied to the syringeal configuration that *α* and *β* jointly set and shares more of their explanatory variance [22]. That RA activity is well predicted by biomechanical control parameters, beyond what the acoustic features capture, is consistent with RA representing vocal output in motor rather than acoustic coordinates, with *α* reflecting the respiratory drive that sets subsyringeal air sac pressure, controlled by RA through brainstem nuclei RAm and PAm, and *β* the drive to syringeal muscles controlled by RA through nXIIts [23]. These conclusions rest on a limited number of birds (n = 5) and should be understood as such.

VIBE opens several avenues for future investigation. For systems neuroscientists, it provides a principled motor representation space against which activity in RA, HVC, and their downstream targets can be directly compared, replacing acoustic features with the variables the motor system actually controls. The continuous recovery of *α* is particularly valuable for studies of respiratory-vocal coupling, where pressure dynamics across syllables and silences carry information about the timing and coordination of motor gestures. For researchers studying vocal learning and plasticity, trajectories through (*α, β*) space provide a compact and physically interpretable description of how the execution of song changes over development or in response to experimental perturbation, complementing acoustics-based measures with a representation grounded in the motor commands being learned. The songbird model system is uniquely suited to studying how a small set of continuously varying control parameters generates a rich acoustic repertoire, and VIBE makes it possible to pursue that question at scale in large datasets and in direct relation to the neural activity that drives it.

## Data Availability

VIBE is openly available at https://github.com/laurenostrowski/VIBE and archived at Zenodo (https://doi.org/10.5281/zenodo.20947855).

## Acknowledgments

The authors thank Lauren Stanwicks and Trevor McPherson for their assistance with data collection and surgical procedures.

## Author Contributions

L.M.O. and T.Q.G. conceived and designed research; L.M.O., J.M.M., and P.T.-M. performed experiments; L.M.O. analyzed data; L.M.O. interpreted results of experiments; L.M.O. prepared figures; L.M.O. drafted manuscript; L.M.O., J.M.M., P.T.-M., B.G.C., and T.Q.G. edited and revised manuscript; L.M.O., J.M.M., P.T.-M., B.G.C., and T.Q.G. approved final version of manuscript.

## References

1. Doupe AJ, Kuhl PK. Birdsong and human speech: Common themes and mechanisms. Annu Rev Neurosci. 1999;22:567–631.

2. Brainard MS, Doupe AJ. What songbirds teach us about learning. Nature. 2002;417(6886):351–8.

3. Vicario DS. Organization of the zebra finch song control system: functional organization of outputs from nucleus robustus archistriatalis. J Comp Neurol. 1991;309(4):486–94.

4. Wild JM. Descending projections of the songbird nucleus robustus archistriatalis. J Comp Neurol. 1993;338(2):225–41.

5. Amador A, Perl YS, Mindlin GB, Margoliash D. Elemental gesture dynamics are encoded by song premotor cortical neurons. Nature. 2013;495(7439):59–64.

6. Arneodo EM, Chen S, Brown DE, Gilja V, Gentner TQ. Neurally driven synthesis of learned, complex vocalizations. Curr Biol. 2021;31(15):3419–25.

7. Gardner T, Cecchi G, Magnasco M, Laje R, Mindlin GB. Simple motor gestures for birdsongs. Phys Rev Lett. 2001;87(20):208101.

8. Laje R, Gardner TJ, Mindlin GB. Neuromuscular control of vocalizations in birdsong: a model. Phys Rev E. 2002;65(5):051921.

9. Arneodo EM, Mindlin GB. Source-tract coupling in birdsong production. Phys Rev E. 2009;79(6 Pt 1):061921.

10. Perl YS, Arneodo EM, Amador A, Mindlin GB. Nonlinear dynamics and the synthesis of Zebra finch song. Int J Bifurc Chaos. 2012;22(10):1250235.

11. Perl YS, Arneodo EM, Amador A, Goller F, Mindlin GB. Reconstruction of physiological instructions from Zebra finch song. Phys Rev E. 2011;84(5 Pt 1):051909.

12. Amador A, Mindlin GB. Beyond harmonic sounds in a simple model for birdsong production. Chaos. 2008;18(4):043123.

13. Sitt JD, Amador A, Goller F, Mindlin GB. Dynamical origin of spectrally rich vocalizations in birdsong. Phys Rev E. 2008;78(1 Pt 1):011905.

14. Boari S, Perl YS, Amador A, Margoliash D, Mindlin GB. Automatic reconstruction of physiological gestures used in a model of birdsong production. J Neurophysiol. 2015;114(5):2912–22.

15. Titze IR. The physics of small-amplitude oscillation of the vocal folds. J Acoust Soc Am. 1988;83(4):1536–52.

16. Sainburg T, Thielk M, Gentner TQ. Finding, visualizing, and quantifying latent structure across diverse animal vocal repertoires. PLoS Comput Biol. 2020;16(10):e1008228.

17. Byrd RH, Lu P, Nocedal J, Zhu C. A limited memory algorithm for bound constrained optimization. SIAM J Sci Comput. 1995;16(5):1190–208.

18. Goller F, Cooper BG. Peripheral motor dynamics of song production in the zebra finch. Ann NY Acad Sci. 2004;1016:130–52.

19. Tostado-Marcos P, Arneodo EM, Ostrowski L, Brown DE, Perez XA, Kadwory A, et al. Population dynamics in songbird RA and HVC during learned motor-vocal behavior. J Neurosci. 2026;46(20):e0580252026.

20. Vicario DS. A new brain stem pathway for vocal control in the zebra finch song system. Neuroreport. 1993;4(7):983–6.

21. Rubner Y, Tomasi C, Guibas LJ. The earth mover’s distance as a metric for image retrieval. Int J Comput Vis. 2000;40(2):99–121.

22. Amador A, Margoliash D. A mechanism for frequency modulation in songbirds shared with humans. J Neurosci. 2013;33(27):11136–44.

23. Schmidt MF, Goller F. Breathtaking songs: coordinating the neural circuits for breathing and singing. Physiology. 2016;31(6):442–51.

